# Interaction of yeast Vps13 with the calcium binding proteins Cdc31 and calmodulin

**DOI:** 10.64898/2026.11.21.768306

**Authors:** Nilufer Dilmen, Jae-Sook Park, Raymond T. Suhandynata, Aaron M. Neiman

**Author notes:** Corresponding author: Room 332 Life Sciences, Stony Brook University, 11794-5215. Current address: Department of Pathology and Skaggs School of Pharmacy and Pharmaceutical Sciences, University of California San Diego, La Jolla, CA.

## Abstract

Vps13 is a member of the bridge-like lipid transfer protein superfamily involved in transport of lipids between different intracellular membranes. In yeast, deletion of *VPS13* exhibits several distinct phenotypes due to preventing interorganelle lipid transfer between many different organelles. Vps13 shuttles lipids through a long hydrophobic channel, but how this activity is regulated in the cell is not yet understood. Studies in yeast and human cells have implicated small calcium-binding proteins, Cdc31 in yeast and calmodulin in mammalian cells, as potential regulators of Vps13 activity. This report identifies the binding site on Vps13 for Cdc31, the yeast ortholog of centrin. The yeast calmodulin protein Cmd1 binds Vps13 through the same site and separate Cmd1-Vps13 and Cdc31-Vps13 complexes co-exist *in vivo*. Association of Cmd1 with Vps13, but not Cdc31, is sensitive to the presence of Ca^2+^. A mutant of *VPS13* that is unable to interact with either Cdc31 or Cmd1 is specifically defective in mitochondrial homeostasis and endoplasmic-reticulum (ER)-phagy, indicating that Vps13 binding to Cdc31 and/or Cmd1 is important for a subset of *VPS13* functions.

## Introduction

Bridge-like lipid transfer proteins (BLTPs) are a superfamily of proteins that mediate the bulk transfer of lipids between intracellular membranes [1, 2]. BLTPs are large rod-shaped proteins characterized by repeated copies of a structural motif, the repeating beta groove (RBG) domain [3]. An RBG is a five stranded antiparallel β-sheet with a bend in the middle of the sheet creating concave and convex faces. Multiple RBG repeats within the protein stack on each other so that the concave faces form a long hydrophobic groove that runs the length of the extended rod creating a channel for lipid transfer [3]. Each end of the channel associates with a different organelle at membrane contact sites, allowing lipids to move between the two membranes [4]. Mutations in different human BLTPs lead to a variety of disorders, such as Neuroacanthocytosis, Cohen Syndrome, Parkinson’s Disease, and a cerebellar ataxia, highlighting the critical importance of this class of transporters [5-8]. How lipid transport between organelles by BLTPs is regulated within the cell is not yet understood. One emerging theme is that partner proteins that bind to the BLTP may control activity of the transport channel itself [9].

Vps13 proteins are one of the best studied families of BLTP [10]. Mammals have four VPS13 paralogs, A, B, C, and D [11]. Different neurological and neurodegenerative disorders are associated with mutations in the genes encoding these paralogs [5-8]. In contrast, the budding yeast, *Saccharomyces cerevisiae*, has a single *VPS13* gene, making it an ideal model for functional studies. Yeast carrying a deletion of *VPS13* are defective in sporulation, vacuolar protein sorting, mitochondrial homeostasis and ER-phagy [12-15]. These different phenotypes reflect the localization of Vps13 to different inter-organelle membrane contact sites [15, 16]. Localization of Vps13 to these different sites is regulated by interactions with organelle-specific adaptor proteins [17-19]. Different adaptors compete with each other for a common binding site in the Vps13 adaptor binding domain, located near the C-terminal end of the transport channel [17]. How the N-terminal end of Vps13 associates with membranes is less clear. In mammalian cells, the ER resident protein VAP binds towards the N-terminal end of Vps13 orthologs to anchor them at the ER, but how yeast Vps13 associates with the ER is not known [20, 21].

In addition to organelle-specific adaptors, another partner of Vps13 is the calcium binding protein Cdc31 [22, 23]. *CDC31* is an essential gene in yeast that encodes a component of the spindle pole body [24]. *CDC31* has also been implicated in coupling of transcription and mRNA export as part of the T-REX2 complex [25-27]. Cdc31, like its mammalian ortholog centrin, is composed of four calcium-binding EF hands, similar to the closely related calmodulin protein [25]. Cdc31 co-purifies with Vps13 from yeast and is necessary for Vps13 to function in an in vitro membrane fusion assay [22]. Moreover, several temperature-sensitive *cdc31* alleles show defects in vacuolar protein sorting, supporting the idea that Vps13 requires Cdc31 for function in vivo [22]. A recent high throughput study identified the yeast calmodulin Cmd1 as a Vps13-interacting protein, suggesting that Vps13 may associate with multiple calcium binding proteins [28]. In human cells, calmodulin has been found to bind to the human Vps13 orthologs VPS13A, VPS13C, and VPS13D, indicating that the interaction between Vps13 and calmodulin is highly conserved [29].

This study identified a binding site for Cdc31 near the N-terminus of Vps13 based on an AlphaFold multimer model of the heterodimer. Biochemical and genetic experiments revealed that Cmd1 binds to the same site as Cdc31 and that the interaction between Vps13 and these calcium-binding proteins regulates a subset of *VPS13* functions in yeast.

## Results

### Cdc31 binds near the N-terminus of Vps13

AlphaFold Multimer was used to produce a putative structure of the Vps13-Cdc31 heterodimer [30](Figure 1A). The algorithm predicted that Cdc31 interacts with Vps13 primarily along a helix (aa 340 to 361 of Vps13) that lies in a loop between the second and third RBG repeats near the N-terminal end of the transport channel (Figure 1A). AlphaFold produced multiple models of the complex, all of which placed Cdc31 at the same location on Vps13, though with varying postures and hydrogen bonding patterns. The model with the highest confidence indicated an extensive contact surface between Vps13 and Cdc31 stabilized by multiple hydrogen bonds (Figure 1A and 1B).

**Figure 1.**
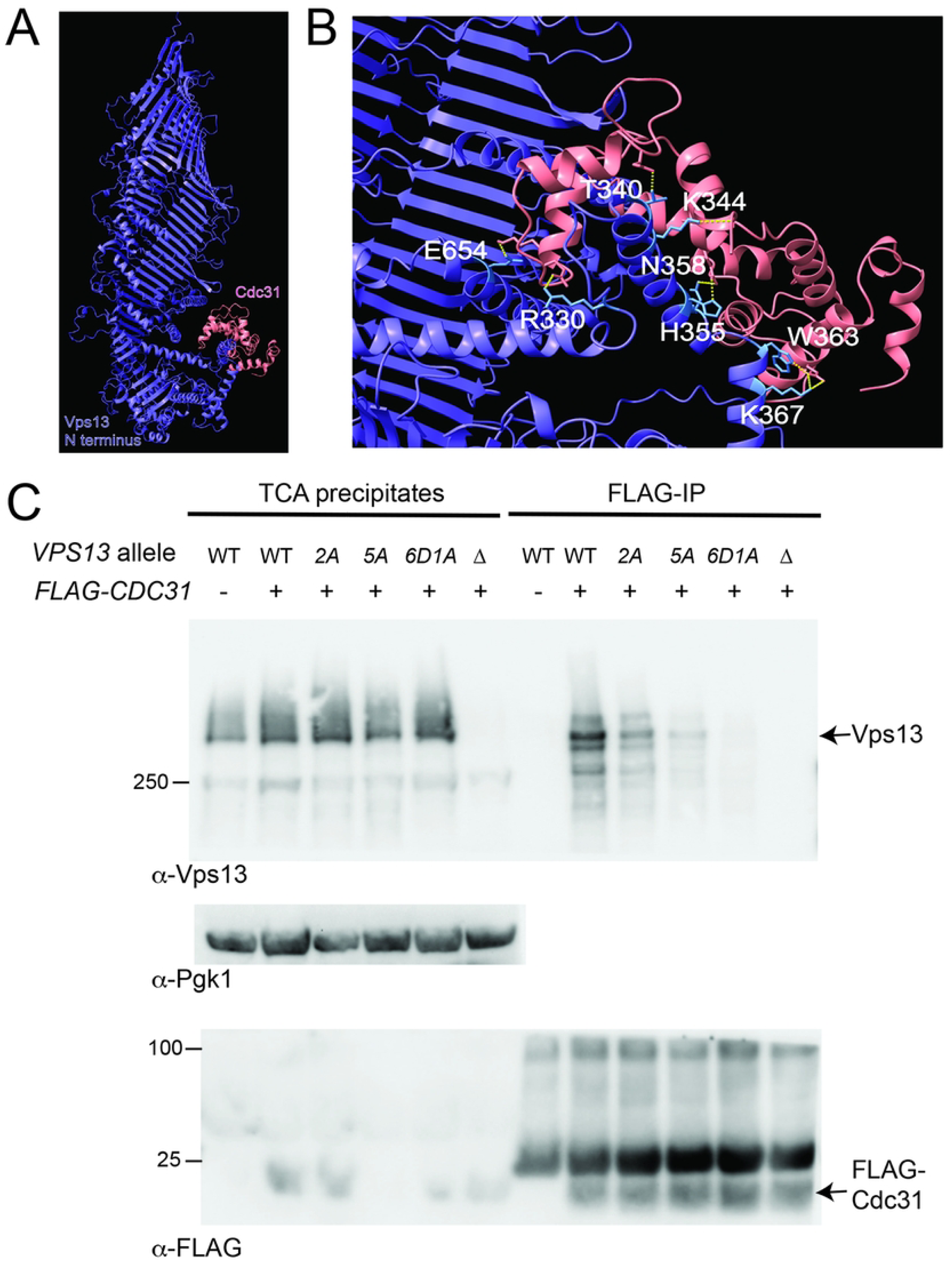
Cdc31 binds to a region near the N-terminal end of Vps13. A) AlphaFold multimer model of the N-terminal half (aa 1-1800) of Vps13 (purple) with Cdc31 (pink). B) Close-up view of the model with Vps13 residues predicted to form hydrogen bonds with Cdc31 highlighted in blue. Hydrogen bonds between the interacting side chains are indicated by yellow dashed lines. C) Point mutations in the predicted hydrogen bonding residues disrupt the Vps13 – Cdc31 interaction. TCA precipitates or α-FLAG immunoprecipitates of WT (BY4741), *vps13-2A* (NDY8), *vps13-5A* (NDY11), *vps13-6D1A* (NDY12) or *<u>vps13</u>*Δ (K01) strains carrying pTEP36 expressing FLAG-Cdc31 were examined by Western blot using α-Vps13 (top panel), α-Pgk1 as a loading control (middle panel), or α-FLAG antibodies. Numbers at left indicate the position of size standards (kDa). Blots shown are representative of four experiments.

To test this model, Vps13 residues that were predicted to form hydrogen bonds with Cdc31 were mutated and the effects of those mutations on the co-precipitation of Vps13 with FLAG-tagged Cdc31 were examined (Figure 1C). Levels of both Vps13 and the FLAG-Cdc31 protein were too low to be detected in the soluble extract, so expression was first checked by whole cell TCA precipitates (Figure 1C). Substitution of Vps13 amino acids R330 and E654 with alanine (*vps13-2A*) partially reduced the co-immunoprecipitation (IP) with FLAG-Cdc31 (Figure 1C). The reduction was exacerbated when residues R330, Y345, H355, N358, and E654 were all mutated to alanine (*vps13-5A*). Changing an amino acid to alanine is not as severe as reversing the charge. Therefore four of the five amino acids in the *vps13-5A* mutant, as well as K344 and K367, were changed instead to the negatively charged aspartate to create the *vps13-6D1A* allele. Vps13^6D1A^ failed to co-precipitate with FLAG-Cdc31 (Figure 1C). Importantly, the Vps13^6D1A^ protein was present at wild-type levels, indicating that the mutations interfere with Cdc31 binding without destabilizing the protein (Figure 1C). These results indicate that this region of Vps13 is the binding site for Cdc31. Interestingly, in mammalian cells, calmodulin binds to the corresponding helix in VPS13C [29].

### Cmd1 binds to the same region of Vps13 as Cdc31

To seek additional Vps13 partner proteins, Vps13-GFP was immunoprecipitated using an α-GFP nanobody and co-precipitating proteins were identified by mass spectrometry. As a specificity control, precipitates from cells expressing a GFP-Spo20^51-91^ fusion were also analyzed [31]. Precipitations were performed from sporulating cells undergoing meiosis II, when Vps13 is predominantly localized to the prospore membrane [14]. To enrich for cells in meiosis II, cells were synchronized in meiosis using an estradiol-inducible *NDT80* system [32]. Cells were incubated in SPO medium for six hours without estradiol, during which time they accumulate in meiotic prophase, and then released into the meiotic divisions by addition of estradiol. The cultures were harvested and IPs performed three hours after estradiol addition when most cells have reached meiosis II and prospore membranes are present [33, 34]. Vps13-GFP interacting proteins were defined as those identified by at least two unique peptides specifically in the Vps13-GFP precipitates that provided greater than 10% coverage of the protein (Table 1)(complete list of all peptides identified is in Supplemental File 1).

**Table 1.** Vps13 co-precipitating proteins identified by mass spectrometry.

| Protein | Percent coverage | # unique peptides |
|---|---|---|
| Cmd1 | 16.3 | 2 |
| Bmh2/Bmh1 | 33.7 | 9 |
| Spo71 | 10.3 | 12 |
| Hsp12 | 27.5 | 3 |
| Pln1 | 21.5 | 5 |
| Fpr3 | 13.2 | 3 |

The sporulation-specific adaptor of Vps13, Spo71, was observed in the Vps13-GFP IP but not the adaptor proteins Ypt35 or Mcp1, which target Vps13 to the vacuole and mitochondria, respectively, during mitotic growth [17, 18]. These results support the assumption that Vps13-GFP was primarily localized at the prospore membranes at the time the extracts were generated for the IP. Further evidence that the IP is detecting bona fide Vps13-GFP interators was the identification of the calmodulin protein Cmd1, the 14-3-3 proteins Bmh1/2, and the yeast perilipin Pln1 [19, 35, 36]. Cmd1 was previously detected as Vps13 binding protein in a high through put IP experiment [28]. Mammalian orthologs of Cmd1 and Bmh1/2 both interact with VPS13D while the human perilipin ortholog PLN3 binds to VPS13C, indicating that these are highly conserved interactions [29, 37].

While Cdc31 was not detected in the Vps13-GFP immunoprecipitate, the close relationship between Cdc31 and Cmd1, as well as the binding site for calmodulin on VPS13C, suggested that Cmd1 interacts with Vps13 through the same site as Cdc31 [29]. Indeed, AlphaFold multimer modeling places Cmd1 binding on Vps13 in the same location as Cdc31 (Figure 2A). Several of the Vps13 residues predicted by AlphaFold to hydrogen bond to Cdc31 (R330, K344, H355, N358) are also predicted by AlphaFold to form hydrogen bonds with Cmd1. Cmd1 and Vps13 also interact in vegetative cells, as endogenous Vps13 protein co-immunoprecipitated with Cmd1-GFP (Figure 2B). In contrast, Vps13^6D1A^ was not detected in the Cmd1-GFP immunoprecipitate indicating that Cmd1 binds to the same site on the Vps13 protein as Cdc31 (Figure 2B).

**Figure 2.**
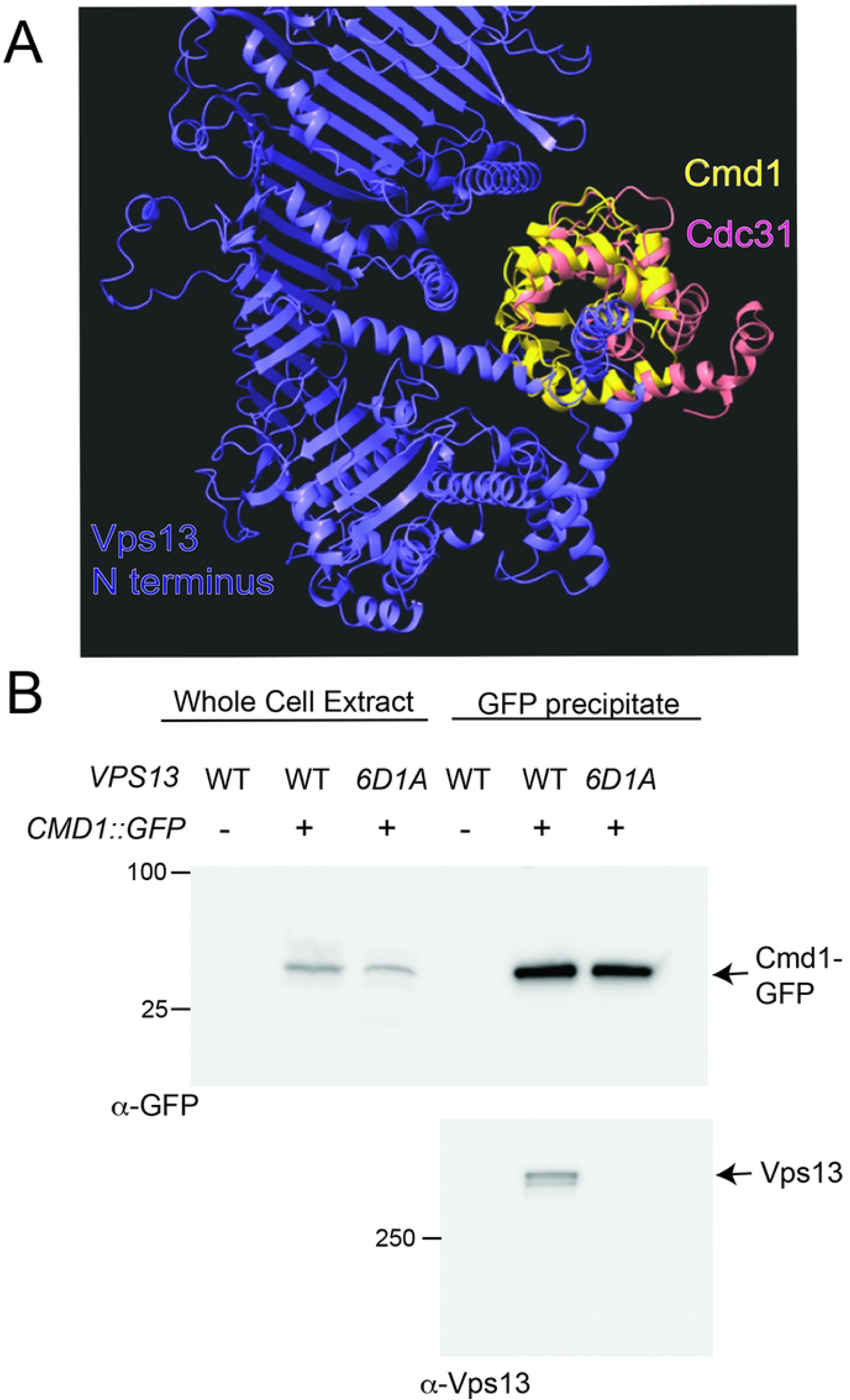
Cmd1 binds to the same site on Vps13 as Cdc31. A) AlphaFold multimer models of the Vps13 N-terminal half (purple) in complex with Cdc31 (pink) and Cmd1 (yellow) were aligned using Chimera X. Blots shown are representative of four experiments. B) Point mutations that disrupt the Vps13-Cdc31 interaction also disrupt Vps13-Cmd1 binding. GFP-Cmd1 was precipitated from extracts of WT (BY4741), *VPS13 CMD1-GFP* (GCY6) or *vps13-6D1A CMD1-GFP* (NDY14) cells and pellets were examined by Western blot using α-GFP (top) or α-Vps13 (bottom) antibodies. Numbers at left indicate the position of size standards (kd). Blots shown are representive of three experiments.

### The Cmd1-Vps13 interaction is sensitive to Ca^2+^

Both Cdc31 and Cmd1 are calcium-binding proteins, raising the possibility that their interactions with Vps13 are affected by the presence or absence of free Ca^2+^. The lysis buffer used in the co-IP experiments contained 5 mM EDTA that is able to chelate both Ca^2+^and Mg^2+^, and should have very low free Ca^2+^ [38]. The Cmd1-GFP/Vps13 co-IP experiments were repeated with the addition of either (1) 5 mM EGTA to chelate any Ca^2+^ not bound by EDTA, (2) 10mM Ca^2+^ to provide free Ca^2+^, or (3) 10 mM Mg^2+^ as a control for any nonspecific effects of a divalent cation. The co-IP of myc-tagged Myo5 with calmodulin was used as a positive control, as this interaction is disrupted by free Ca^2+^ [39]. Addition of EGTA had no effect on the co-IP of myc-tagged Myo5 with Cmd1-GFP, while addition of Ca^2+,^ but not Mg^2+^, disrupted the interaction (Figure 3A). The co-IP of Vps13 with Cmd1-GFP was also unaffected by addition of EGTA but specifically reduced by addition of Ca^2+^ (Figure 3B). By contrast, Vps13 co-precipitated with FLAG-Cdc31 under all conditions (Figure 3C). Thus, although Vps13 binds both Cdc31 and Cmd1 in the same location, the two proteins behave differently with respect to the calcium sensitivity of Vps13 binding.

**Figure 3.**
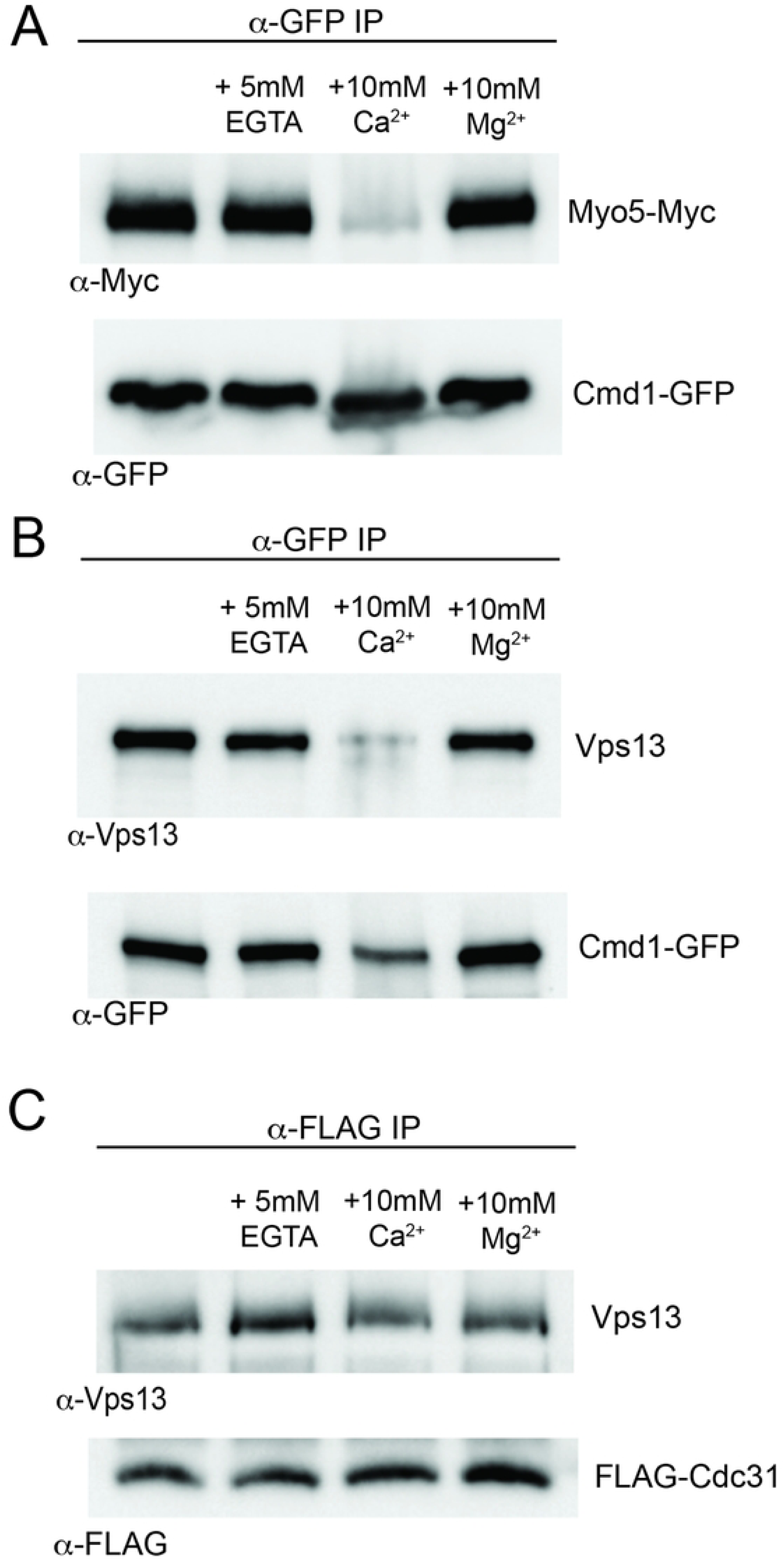
The Cmd1-Vps13 interaction is sensitive to Ca^2+^. A) α-GFP IPs were performed from extracts of GCY6 (*CMD1-GFP*) expressing myc-tagged Myo5 (from pRS-Y2-myc-Myo5) made in standard conditions (5mM EDTA) or in lysis buffer with added EGTA, Ca^2+^ or Mg^2+^, as indicated. The pellets were analyzed by Western blot with α-myc and α-GFP antibodies. B) Strain GCY6 without the myc-MYO5 plasmid was analyzed as in (A) and pellets were probed on a Western blot with α-Vps13 and α-GFP antibodies. C) IPs under the same set of conditions as in (A) were performed using the WT strain (BY4741) carrying a plasmid expressing FLAG-tagged Cdc31 (pTEP36). Pellets tested by Western blot with α-Vps13 and α-FLAG antibodies. Blots shown are representative of two experiments for panel A and four experiments for B and C.

### Cmd1-Vps13 and Cdc31-Vps13 complexes co-exist in the cell

The fact that Cmd1 and Cdc31 bind the same site on Vps13 raises the question of whether distinct Cmd1-Vps13 and Cdc31-Vps13 complexes co-exist in the cell or whether they are mutually exclusive. If the complexes co-exist, then it should be possible to co-immunoprecipitate both complexes from the same extract. To examine this possibility, an extrat from cells containing Cmd1-GFP and either only endogenous Cdc31 or also overexpressed GLAG-Cdc31 was generated from vegetative cells. FLAG-CDC31 was overexpressed in these strains from a high copy plasmid under either the *GPD1* or the *CUP1* promoter. FLAG-Cdc31 was then precipitated with, α-FLAG antibodies and co-IP of Vps13 was observed only in strains carrying the tagged Cdc31 (Figure 4A). The supernatant from the FLAG IP was then incubated with α-GFP nanobodies to precipitate Cmd1-GFP. Vps13 was also observed in the Cmd1-GFP Ips indicating that both complexes were present in the extract (Figure 4B).

**Figure 4.**
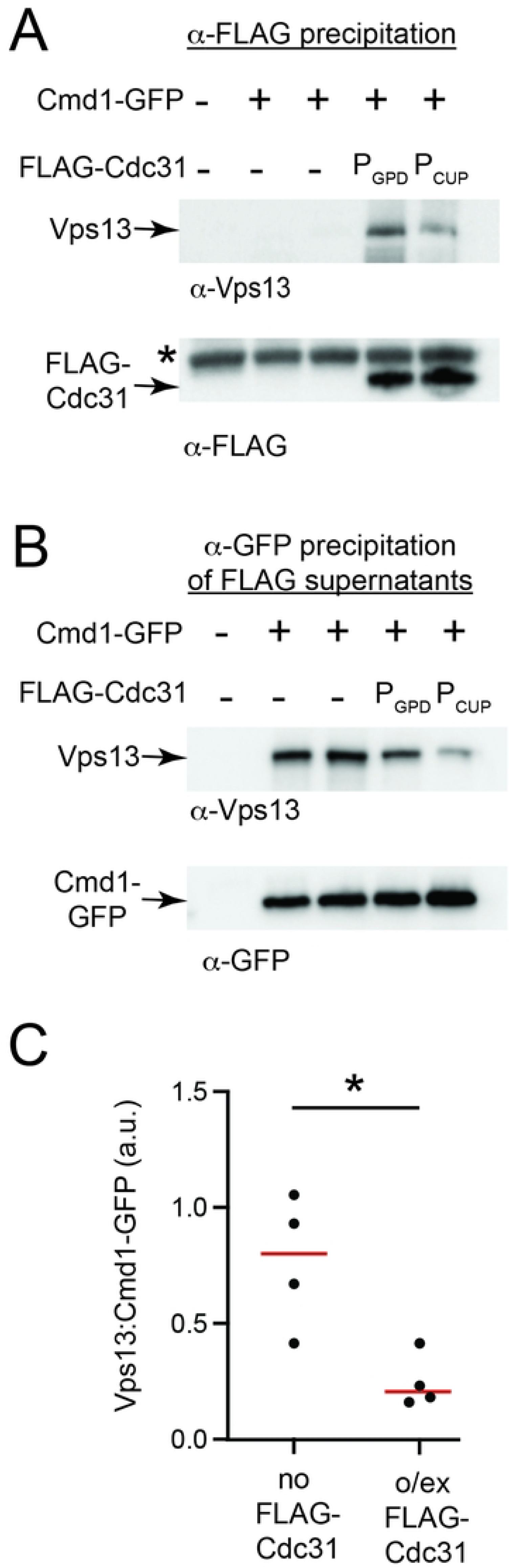
Cdc31 and Cmd1 compete for binding to Vps13. A) Extracts of WT (BY4741) (lane 1), or Cmd1-GFP cells (GCY6) carrying no vector (lane 2), pRS426-P_GPD_ (lane 3), pRS426-P_GPD_-FLAG-Cdc31 (lane 4), or TEP36 (P_CUP1_-FLAG-CDC31; lane 5), were subject to IP using anti-FLAG antibodies. The resulting pellets were separated on SDS-PAGE and examined by Western blot using α-Vps13 or α-FLAG antibodies. Asterisk indicates a cross-reacting band in the anti-FLAG blot. B) The remaining supernatants after anti-FLAG precipitation were incubated with GFP-Trap to precipitate Cmd1-GFP and the resulting pellets examined using α-Vps13 and α-GFP antibodies. C) The amount of Vps13 co-precipitating with Cmd1 was quantified relative to the amount of Cmd1 precipitated in four different samples, either with or without FLAG-Cdc31 overexpression. Red lines indicate the mean. * Indicates the difference is significant at p< 0.03 by Mann-Whitney test. “a.u”, arbitrary units. Blots shown are representative of three experiments.

Interestingly, the amount of Vps13 in the Cmd1-GFP IPs was greater when the FLAG supernatants came from the cells with only the endogenous level of Cdc31 (Figure 4B). Quantification of the ratio of Vps13 to Cmd1-GFP in the precipitates from four biological replicates indicated that overexpression of Cdc31 reduces the amount of Cmd1 bound to Vps13 by about 65%, consistent with the idea that Cdc31 and Cmd1 compete for binding to Vps13 (Figure 4C).

### Mutation of the Cdc31/Cmd1 binding domain in *VPS13* causes defects in mitochondrial homeostasis and ER-phagy

Cdc31 is required for the activity of Vps13 in an *in vitro* membrane fusion assay and some *cdc31-ts* alleles display vacuolar protein sorting defects similar to *vps13Δ* mutants, suggesting that the interaction of Vps13 with Cdc31 is important for function [22]. To explore the role of Vps13 interaction with calcium binding proteins *in vivo*, cells carrying the Cdc31/Cmd1 interaction defective *vps13-6D1A* allele were examined for four different phenotypes seen in *vps13*Δ mutants: sorting of carboxypeptidase Y to the vacuole, sporulation defects, synthetic lethality with *mmm1*Δ (a readout of mitochondrial homeostasis), and defects in autophagy of the peripheral ER [12-15] (Figure 5).

**Figure 5.**
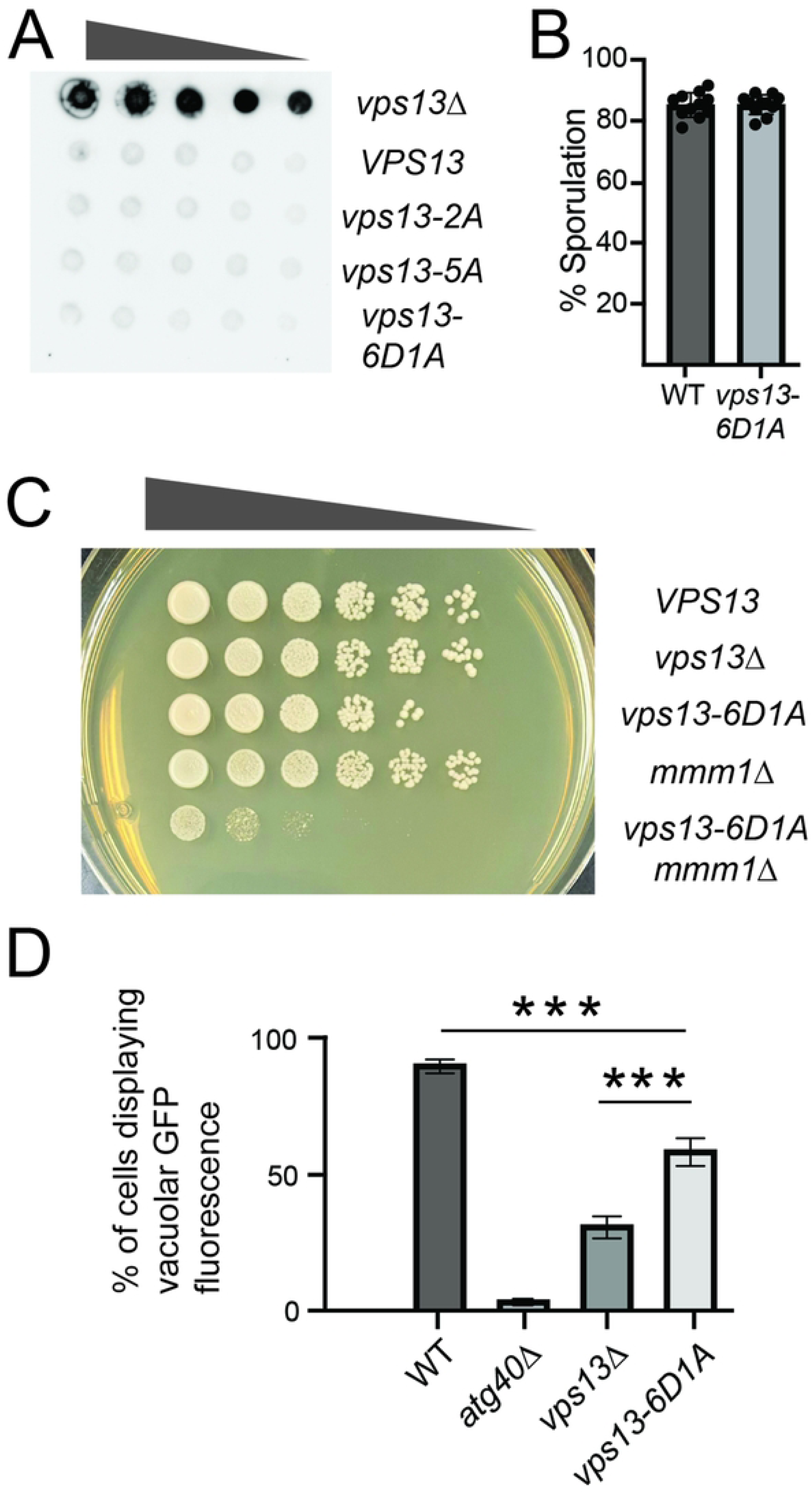
The *vps13-6D1A* mutant displays a synthetic phenotype with *mmm1*Δ and ER-phagy defects. A) *vps13-6D1A* displays no defect in vacuolar sorting. 4-fold serial dilutions of WT (BY4741), *vps13*Δ (K01) *vps13-2A* (NDY8), *vps13-5A* (NDY11), and *vps13-6D1A* were spotted onto SC plates, overlain with a nitrocellulose membrane, and incubated overnight at 30°. The filters were then washed and probed for Carboxypeptidase Y secretion using α-CPY antibodies. B) *vps13-6D1A* displays no sporulation defect. Diploids heterozygous for *vps13*Δ and *VPS13* (NDY22) or *vps13-6D1A* (NDY23) were sporulated at 30° and assessed for spore formation by light microscopy. At least 400 cells were scored in three separate experiments. Bars show one standard deviation. C) *vps13-6D1A* shows defects in mitochondrial function. 10-fold serial dilutions of WT (BY4741), *vps13*Δ (K01), *vps13-6D1A* (NDY12), *mmm1*Δ (K04), and *vps13-6D1A mmm1*Δ (NDY24) cells were spotted onto YPD plates and incubated at 30°C. D) *vps13-6D1A* shows defects in ER phagy. WT (GCY5), *atg40*Δ (JSP845), *vps13*Δ (JSP846) or *vps13-6D1A* (NDY20) strains carrying an *RTN1-GFP* fusion were treated with rapamycin overnight at to induce autophagy and the scored for the presence of GFP fluorescence within the vacuole. At least 10 independent measurements were made for each strain at each temperature. Bars indicate one standard deviation. “***” indicates p<0.001 by Mann-Whitney test.

Defects in vacuolar sorting result in the secretion of proteins that would otherwise be trafficked to the vacuole [12]. A commonly used marker to monitor vacuolar sorting is the vacuolar protease Carboxypeptidase Y (CPY). CPY sorting was assessed by spotting serial dilutions of different strains onto SC medium. A nitrocellulose filter was placed on top of the cells that were then allowed to grow overnight at 30°C or 37°C. The membranes were then removed and probed with α-CPY antibodies to detect secreted, and therefore mis-sorted, CPY [40]. Unexpectedly, given an earlier report of a CPY sorting defect in *cdc31*-ts mutants [22], *vps13-6D1A* cells displayed no obvious defect in CPY sorting (Figure 5A).*VPS13* is required to make spores after cells have been induced to undergo meiosis [14]. Because sporulation occurs in diploids, a *vps13-6D1A* haploid was crossed with a *vps13*Δ haploid and the hemizygous diploid was examined for sporulation efficiency. The *vps13-6D1A* fully complemented *vps13*Δ, producing an equivalent number of asci as the hemizygous wild-type diploid (Figure 5B).

*MMM1* encodes a component of the ERMES complex that functions redundantly with *VPS13* in lipid transfer to the mitochondrion [15, 16]. While *mmm1Δ* or *vps13Δ* haploids are viable, the double mutant is synthetically lethal [15, 16]. A *vps13-6D1A mmm1*Δ strain was constructed by sporulation and dissection of a *trans* heterozygous diploid. The *vps13-6D1A mmm1*Δ haploid cells were grown to stationary phase and spotted onto rich medium. In contrast to either single mutant, the double mutant cells were very slow-growing (Figure 5C). These results indicate that interaction with Cdc31 and/or Cmd1 is important for the function of Vps13 at mitochondria.

Selective autophagy of the endoplasmic reticulum (ER-phagy) is stimulated by nitrogen starvation and *vps13*Δ mutants are specifically defective in ER-phagy of the cortical ER [13]. To assess ER-phagy, *vps13-6D1A* cells expressing the cortical ER marker *RTN1-GFP* were treated overnight with rapamycin to simulate nitrogen starvation. ER-phagy of the cortical ER can then be monitored in the fluorescence microscope by the appearance of GFP fluorescence in the vacuolar lumen [41]. As reported previously, *vps13*Δ cells displayed defects in ER-phagy after treatment with rapamycin, similar to but slightly less severe than, mutants in the ER-phagy receptor *ATG40* [13]. Cells carrying *vps13-6D1A* displayed a modest but significant ER-phagy defect, intermediate between *vps13*Δ and wild-type (Figure 5D). Therefore, similar to mitochondrial homeostasis, the interaction of Vps13 with Cmd1 and/or Cdc31 is also important but not essential for its role in ER-phagy. Together these observations indicate that association with Cmd1 and/or Cdc31 is important for Vps13 function in some but not all contexts.

## Discussion

A connection between Vps13 proteins and calcium binding proteins was previously established in both mammals and yeast. In mammalian cells, calmodulin binds to three of the four Vps13 paralogs [29]. In yeast, Vps13 copurifies with Cdc31 and high throughput mass spectrometry detected an interaction between Cmd1 and Vps13 [22, 23, 28]. We report here that Vps13 binds to both Cdc31 and Cmd1 along the linker between the second and third RBG repeats, near the amino terminal end of the lipid transfer channel. The cryo-EM structure of a VPS13C-calmodulin complex revealed that calmodulin binds in the corresponding region of mammalian VPS13C, reinforcing the conserved nature of this interaction [29]. Cdc31 is also highly conserved, with three orthologs (Centrin-1, -2, and -3) in human cells. Whether any of the mammalian VPS13 proteins associate with any of the centrin proteins has not been reported, but given the high conservation of the proteins involved it would not be surprising if the mammalian VPS13 proteins, similar to yeast Vps13, interact with both centrin and calmodulin.

Despite this strong conservation, however, the impact of disrupting the interaction between Vps13 and Cdc31/Calmodulin is quite different in yeast and mammalian cells. Loss of the interaction in yeast affects only a subset of Vps13 functions and the mutant Vps13 protein is stable. By contrast, mutations in VPS13C that disrupt calmodulin binding alter the localization and stability of the protein [29]. Further, one such mutation *VPS13C-W395C*, was originally identified in Parkinson’s patients, suggesting that the mutation causes strong functional defects *in vivo* [42]. Thus, the interaction with calmodulin seems to be a more significant feature of VPS13C function than it is for yeast Vps13. VPS13A and VPS13D have also been shown to bind to calmodulin. It will be interesting to learn if mutations disrupting calmodulin binding in these orthologs behave more like the mutations in VPS13C or the yeast Vps13.

The *vps13-6D1A* allele that disrupts Cdc31/Cmd1 binding showed defects in mitochondrial homeostasis and in ER-phagy, but no defects in CPY sorting or sporulation. The lack of CPY sorting phenotype was unexpected given earlier studies implicating Cdc31 in this Vps13 function [22]. The reason for the discrepancy is not clear. One possibility is that the reported importance of Cdc31 in CPY sorting reflects a requirement for Cdc31 to interact with some other factor involved in sorting besides Vps13. Alternatively, it is possible that the mutations in the *vps13-6D1A* allele are sufficient to disrupt the co-precipitation with Cdc31 under our *in vitro* conditions but do not completely block the interaction of the proteins *in vivo*. As the effects of *VPS13* on ER-phagy are genetically separable from its mitochondrial role, our results nonetheless indicate that the interaction with Cdc31/Cmd1 is important for multiple functions of Vps13 [41].

Though Cdc31 and Cmd1 compete for binding to the same region of Vps13, the effects of Ca^2+^ on their interaction with Vps13 differs. The Cdc31 association with Vps13 was not sensitive to the presence of free Ca^2+^, while the interaction of Cmd1 with Vps13 was lost when free Ca^2+^ was present. This latter result contrasts with the interaction between calmodulin and VPS13C in mammalian cells, where the effect of Ca^2+^ on the interaction is more subtle [29]. Though the presence of free Ca^2+^ does not disrupt the calmodulin-VPS13C co-precipitation, hydrogen-deuterium mass spectrometry analysis indicates that the conformation of calmodulin and VPS13C is altered when Ca^2+^ is present [29]. Whether these *in vitro* differences occur *in vivo* is not clear, but these observations suggest that the interaction between calmodulin and VPS13C may be more stable than that between the yeast orthologs.

The different sensitivities to Ca^2+^ displayed by Cmd1 and Cdc31 raises the possibility that Ca^2+^ influx could regulate a switch between Vps13-Cmd1 and Vps13-Cdc31. That is, introduction of Ca^2+^ could disrupt Vps13-Cmd1 complexes, thereby releasing Vps13 to now bind to Cdc31. Furthermore, the competition between Cmd1 and Cdc31 binding to a site near the N- terminus of Vps13 is similar to the situation with adaptor proteins, which compete for binding to a site near the C-terminal end of the transfer channel [17]. In this case, the different adaptors anchor Vps13 to different organelles and promote distinct intracellular roles for Vps13. Thus, by analogy, it is possible that Vps13-Cdc31 and Vps13-Cmd1 complexes have different functions and that Ca^2+^ could regulate a switch between these functions. Alleles of *VPS13* that specifically disrupt binding of Cdc31 or Cmd1 would be important tools for testing this possibility. As the *vps13-6D1A* allele abrogates binding of both proteins, whether these two complexes have distinct functions remains an open question.

## Methods

### Yeast Strains and Culture

Unless otherwise mentioned, standard yeast media and genetic techniques were used [43]. Yeast strains used are listed in Table 2. All the *VPS13* point mutations were introduced into the genome using a two-step CRISPR/Cas9 strategy as described in [15]. To create a targeted double strand break within the *VPS13* gene, a *SkHIS3MX6* cassette was first introduced by PCR-mediated integration into a region of the *VPS13* open reading frame where the mutation was to be made [44]. The *vps13::SkHIS3MX6* haploids were then cotransformed with pRS425-Cas9-SkHIS3-381, which carries the CAS9 gene as well as a sequence encoding a guide RNA that targets a double strand break to *SkHIS3* [45], and a ∼75 residue oligonucleotide encoding the desired codon changes as a healing fragment to allow the double strand break to be repaired and the mutation(s) incorporated by homologous recombination. After selection for transformants carrying the plasmid, colonies were screened for the loss of the *HIS3* marker, and the presence of the mutations was confirmed by sequencing a PCR product spanning the modified region using Plasmidsaurus (https://plasmidsaurus.com). Using this method, *HIS3MX6* was inserted in place of codon 301 in the *VPS13* coding sequence (strain JSP921) and then *VPS13-R330* was created by removal and repair of the *SkHIS3MX6* cassette (strain NDY1). The *SkHIS3MX6* cassette was then used to replace codon 654 in NDY1 to generate NDY7 and replacement of this insertion yielded NDY8 (*VPS13-2A*). Again, the *SkHIS3MX6* cassette was used to replace codons 345 to 348 of VPS13 in NDY8, creating NDY10 and replacement in this strain with different rescue oligos produced NDY11 (*VPS13-5A*) and NDY12 (*vps13-6D1A*).

**Table 2.**
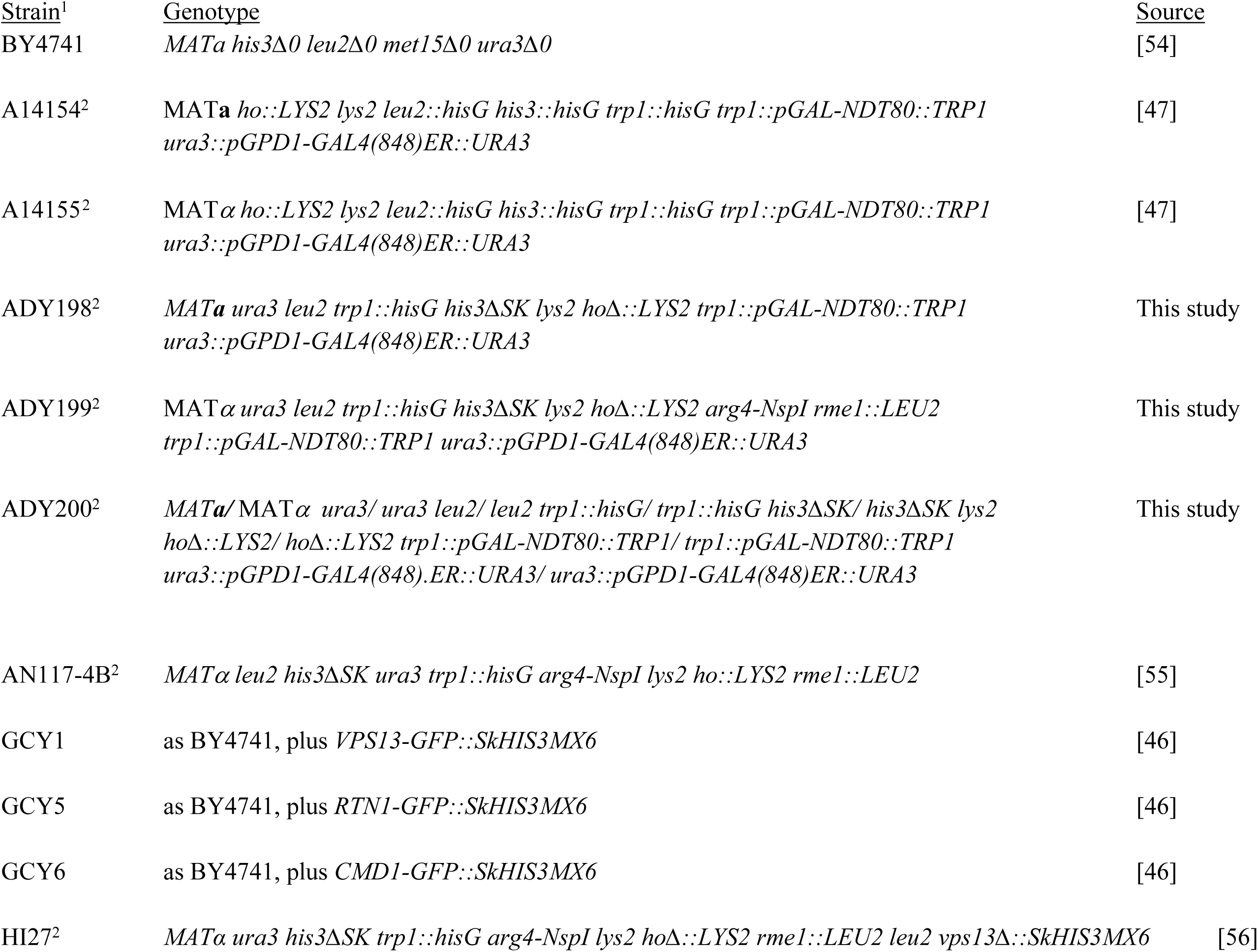
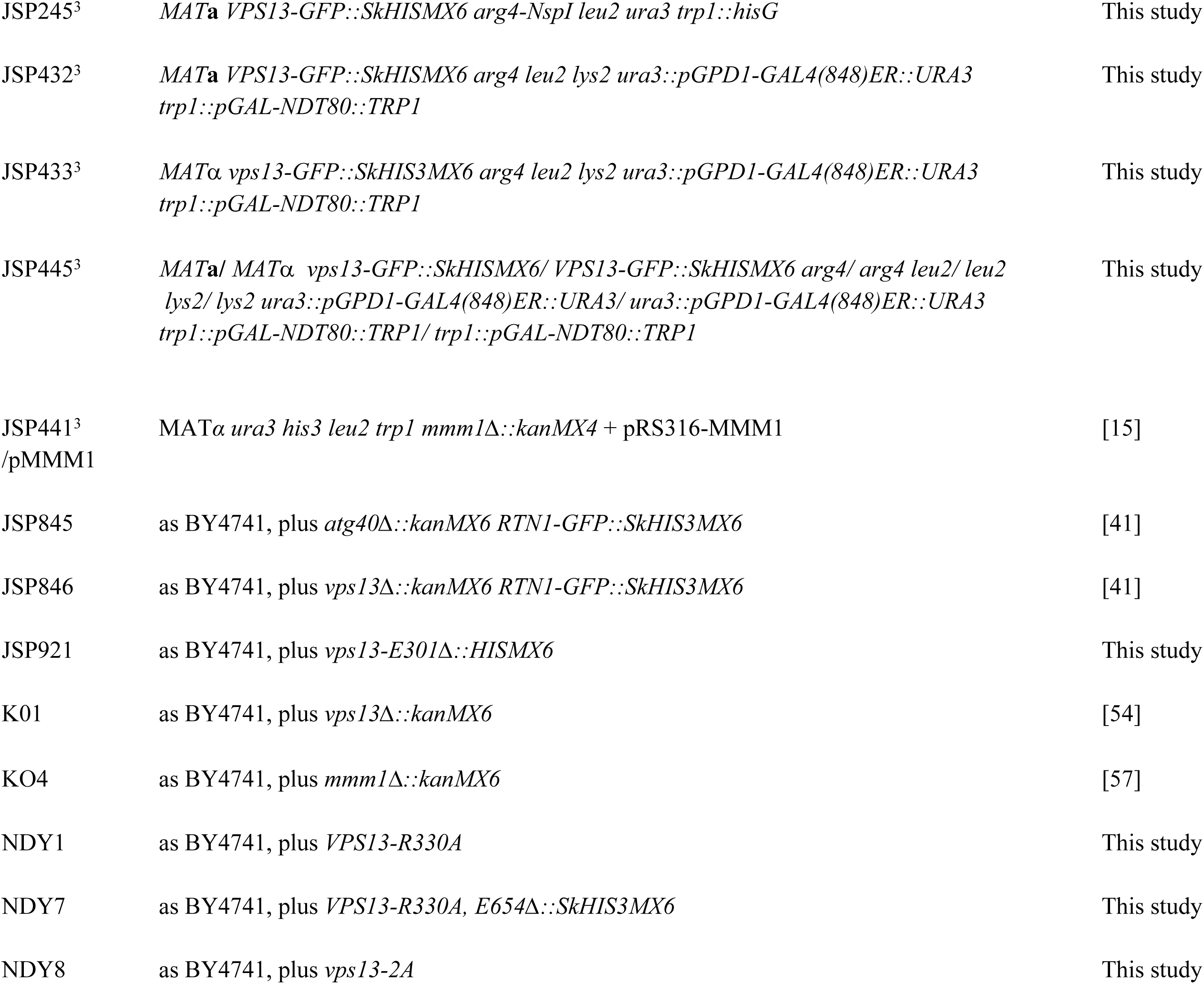
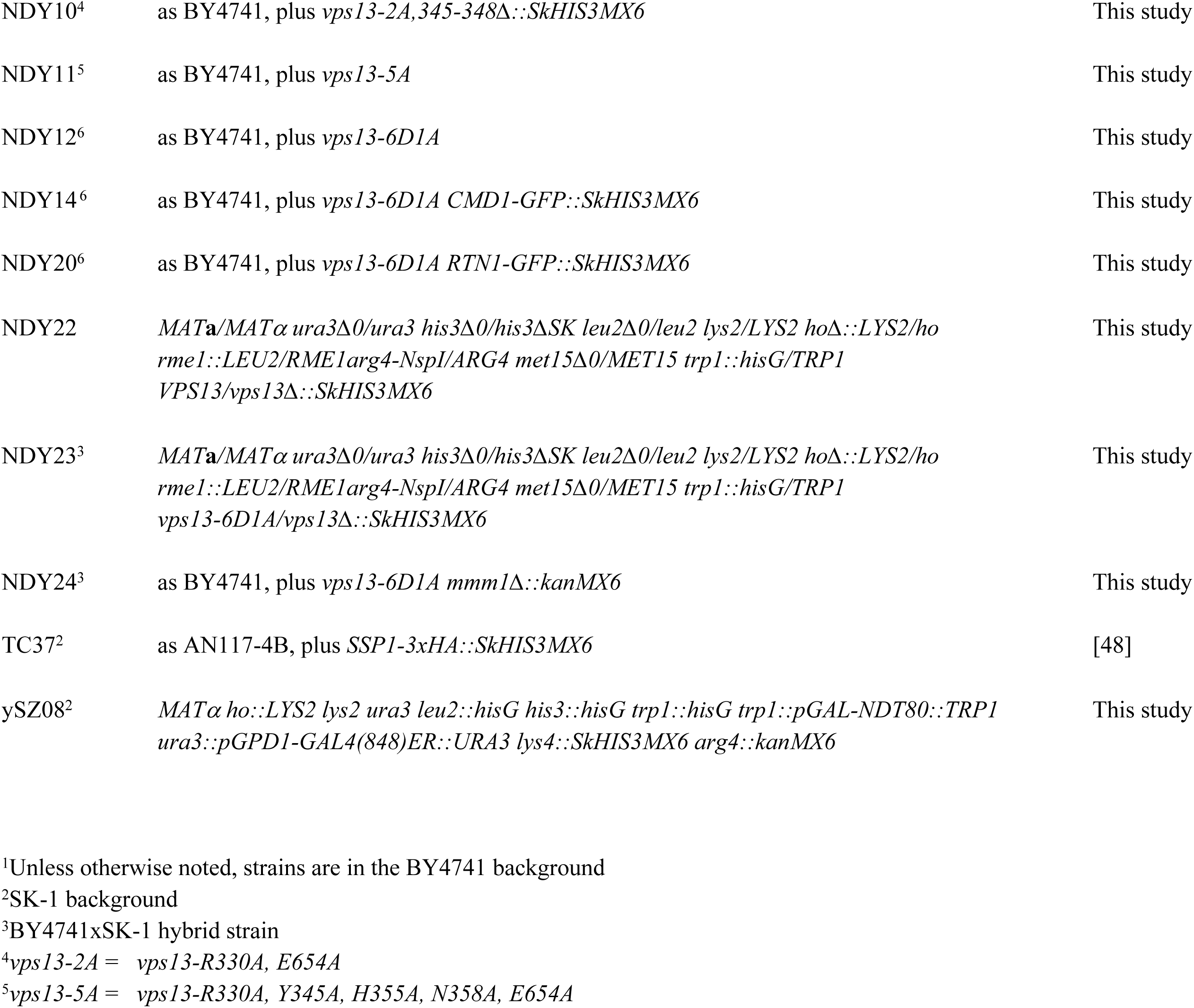
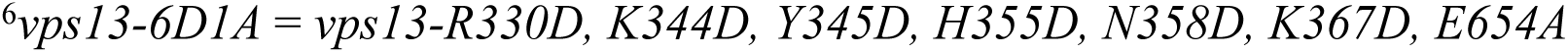
Strains used in this study.

| Strain<sup>1</sup> | Genotype | Source |
|---|---|---|
| BY4741 | *MATa his3Δ0 leu2Δ0 met15Δ0 ura3Δ0* | [54] |
| A14154<sup>2</sup> | MAT**a** *ho::LYS2 lys2 leu2::hisG his3::hisG trp1::hisG trp1::pGAL-NDT80::TRP1 ura3::pGPD1-GAL4(848)ER::URA3* | [47] |
| A14155<sup>2</sup> | MAT*α ho::LYS2 lys2 leu2::hisG his3::hisG trp1::hisG trp1::pGAL-NDT80::TRP1 ura3::pGPD1-GAL4(848)ER::URA3* | [47] |
| ADY198<sup>2</sup> | *MAT**a** ura3 leu2 trp1::hisG his3ΔSK lys2 hoΔ::LYS2 trp1::pGAL-NDT80::TRP1 ura3::pGPD1-GAL4(848)ER::URA3* | This study |
| ADY199<sup>2</sup> | MAT*α ura3 leu2 trp1::hisG his3ΔSK lys2 hoΔ::LYS2 arg4-NspI rme1::LEU2 trp1::pGAL-NDT80::TRP1 ura3::pGPD1-GAL4(848)ER::URA3* | This study |
| ADY200<sup>2</sup> | *MAT**a/*** MAT*α ura3/ ura3 leu2/ leu2 trp1::hisG/ trp1::hisG his3ΔSK/ his3ΔSK lys2 hoΔ::LYS2/ hoΔ::LYS2 trp1::pGAL-NDT80::TRP1/ trp1::pGAL-NDT80::TRP1 ura3::pGPD1-GAL4(848).ER::URA3/ ura3::pGPD1-GAL4(848)ER::URA3* | This study |
| AN117-4B<sup>2</sup> | *MATα leu2 his3ΔSK ura3 trp1::hisG arg4-NspI lys2 ho::LYS2 rme1::LEU2* | [55] |
| GCY1 | as BY4741, plus *VPS13-GFP::SkHIS3MX6* | [46] |
| GCY5 | as BY4741, plus *RTN1-GFP::SkHIS3MX6* | [46] |
| GCY6 | as BY4741, plus *CMD1-GFP::SkHIS3MX6* | [46] |
| HI27<sup>2</sup> | *MATα ura3 his3ΔSK trp1::hisG arg4-NspI lys2 hoΔ::LYS2 rme1::LEU2 leu2 vps13Δ::SkHIS3MX6* | [56] |
| JSP245<sup>3</sup> | *MAT***a** *VPS13-GFP::SkHISMX6 arg4-NspI leu2 ura3 trp1::hisG* | This study |
| JSP432<sup>3</sup> | *MAT***a** *VPS13-GFP::SkHISMX6 arg4 leu2 lys2 ura3::pGPD1-GAL4(848)ER::URA3 trp1::pGAL-NDT80::TRP1* | This study |
| JSP433<sup>3</sup> | *MATα vps13-GFP::SkHIS3MX6 arg4 leu2 lys2 ura3::pGPD1-GAL4(848)ER::URA3 trp1::pGAL-NDT80::TRP1* | This study |
| JSP445<sup>3</sup> | *MAT***a**/ *MATα vps13-GFP::SkHISMX6/ VPS13-GFP::SkHISMX6 arg4/ arg4 leu2/ leu2 lys2/ lys2 ura3::pGPD1-GAL4(848)ER::URA3/ ura3::pGPD1-GAL4(848)ER::URA3 trp1::pGAL-NDT80::TRP1/ trp1::pGAL-NDT80::TRP1* | This study |
| JSP441<sup>3</sup> /pMMM1 | MAT*α ura3 his3 leu2 trp1 mmm1Δ::kanMX4* + pRS316-MMM1 | [15] |
| JSP845 | as BY4741, plus *atg40Δ::kanMX6 RTN1-GFP::SkHIS3MX6* | [41] |
| JSP846 | as BY4741, plus *vps13Δ::kanMX6 RTN1-GFP::SkHIS3MX6* | [41] |
| JSP921 | as BY4741, plus *vps13-E301Δ::HISMX6* | This study |
| K01 | as BY4741, plus *vps13Δ::kanMX6* | [54] |
| KO4 | as BY4741, plus *mmm1Δ::kanMX6* | [57] |
| NDY1 | as BY4741, plus *VPS13-R330A* | This study |
| NDY7 | as BY4741, plus *VPS13-R330A, E654Δ::SkHIS3MX6* | This study |
| NDY8 | as BY4741, plus *vps13-2A* | This study |
| NDY10<sup>4</sup> | as BY4741, plus *vps13-2A,345-348Δ::SkHIS3MX6* | This study |
| NDY11<sup>5</sup> | as BY4741, plus *vps13-5A* | This study |
| NDY12<sup>6</sup> | as BY4741, plus *vps13-6D1A* | This study |
| NDY14<sup>6</sup> | as BY4741, plus *vps13-6D1A CMD1-GFP::SkHIS3MX6* | This study |
| NDY20<sup>6</sup> | as BY4741, plus *vps13-6D1A RTN1-GFP::SkHIS3MX6* | This study |
| NDY22 | *MAT***a***/MATα ura3Δ0/ura3 his3Δ0/his3ΔSK leu2Δ0/leu2 lys2/LYS2 hoΔ::LYS2/ho rme1::LEU2/RME1arg4-NspI/ARG4 met15Δ0/MET15 trp1::hisG/TRP1 VPS13/vps13Δ::SkHIS3MX6* | This study |
| NDY23<sup>3</sup> | *MAT***a***/MATα ura3Δ0/ura3 his3Δ0/his3ΔSK leu2Δ0/leu2 lys2/LYS2 hoΔ::LYS2/ho rme1::LEU2/RME1arg4-NspI/ARG4 met15Δ0/MET15 trp1::hisG/TRP1 vps13-6D1A/vps13Δ::SkHIS3MX6* | This study |
| NDY24<sup>3</sup> | as BY4741, plus *vps13-6D1A mmm1Δ::kanMX6* | This study |
| TC37<sup>2</sup> | as AN117-4B, plus *SSP1-3xHA::SkHIS3MX6* | [48] |
| ySZ08<sup>2</sup> | *MATα ho::LYS2 lys2 ura3 leu2::hisG his3::hisG trp1::hisG trp1::pGAL-NDT80::TRP1 ura3::pGPD1-GAL4(848)ER::URA3 lys4::SkHIS3MX6 arg4::kanMX6* | This study |

<sup>1</sup>Unless otherwise noted, strains are in the BY4741 background
<sup>2</sup>SK-1 background
<sup>3</sup>BY4741xSK-1 hybrid strain
<sup>4</sup>*vps13-2A = vps13-R330A, E654A*
<sup>5</sup>*vps13-5A = vps13-R330A, Y345A, H355A, N358A, E654A*
<sup>6</sup>*vps13-6D1A = vps13-R330D, K344D, Y345D, H355D, N358D, K367D, E654A*

The strain for mass spectrometry identification of Vps13 binding proteins, JSP445, was constructed in several steps. The *VPS13-GFP* allele was obtained from the GFP collection (provided by H. Madhani) in the BY4741 background [46]. To move this allele into the higher sporulating SK1 background, GCY1 (*VPS13-GFP*) was crossed to the SK1 haploid AN117-4B, the diploid sporulated and dissected and a haploid segregant containing *VPS13-GFP* was named JSP245. To introduce the *VPS13-GFP* into an inducible *NDT80* strain background, JSP245 was crossed to ySZ08, which contains the *GAL4-ER* and *P_GAL1_-NDT80* genes. Dissection of the resulting diploid generated two haploid segregants of opposite mating type that contained *VPS13-GFP*, *GAL4-ER* and *P_GAL1_-NDT80*. These haploids were mated to create JSP445. As a negative control for the IP, a *VPS13 GAL4-ER P_GAL1_-NDT80* diploid containing *GFP* was created by crossing the SK1 strain A14154 that contains *GAL4-ER* and *P_GAL1_-NDT80*[47] to a second SK1 haploid, TC37 [48] and dissecting tetrads from the sporulated diploid. Spore colonies of opposite mating type containing *GAL4-ER* and *P_GAL1_-NDT80* (ADY198 and ADY199) were mated to make the homozygous diploid ADY200. The *LEU2 GFP-SPO21^51-91^* plasmid, pRS425-GFP-Spo20^51-91^ was then transformed into ADY200, selecting for Leu^+^ cells.

To construct strains for ER-phagy assays and for Cmd1 co-precipitation assays the *RTN1-GFP::SkHIS3MX6* or *CMD1-GFP::SkHIS3MX6* alleles were amplified by PCR from genomic DNA of GCY5 and GCY6, respectively, and independently introduced into the *vps13-6D1A* haploid (NDY12) by transformation and selection of His^+^ transformants. The resulting haploids are NDY20 (*RTN1::GFP vps13-6D1A*) and NDY14 (*CMD1-GFP vps13-6D1A*).

To make the wild-type and *vps13-6D1A* hemizygous diploids for measuring sporulation, BY4741 or NDY12 (*VPS13* and *vps13-6D1A*, respectively, were mated to HI27 and then diploid cells selected SD-His-Trp medium creating NDY22 and NDY23. To generate the *vps13-6D1A mmm1*Δ strain NDY24, the *mmm1*Δ strain JSP441/pMMM1 was crossed to NDY12 (*vps13-6D1A*) and diploids were selected for on SD-Ura-Trp plates. Diploids were sporulated on Spo medium at 30 °C for six days. Tetrads were dissected, and individual spores were grown on YPD plates for 2 days. The *VPS13* locus was amplified by PCR and sequenced from multiple G418- resistant segregants (*mmm1ΔkanMX6*) that were Ura^-^ (lacking pRS316-MMM1) to identify segregants carrying *vps13-6D1A*. NDY24 is one such segregant.

### Plasmids

The plasmids used are listed in Table 3. To construct a high copy number *URA3* plasmid carrying the *FLAG-CDC31* allele under the control of the constitutive *GPD1* promoter (pRS426-GPD-FLAG-CDC31), Gibson Assembly was performed using the HiFi DNA Assembly kit (New England Biolabs cat. M5520AA). To construct pRS426-GPD-FLAG-CDC31, plasmid TEP36 [49] was used as a template for PCR amplification of the *FLAG-CDC31* insert. A fragment containing *FLAG-CDC31* was generated using PCR and the template plasmid TEP36 [49]. The pRS426-GPD vector [50] was next linearized by digestion with BamHI and HindIII. The amplified *FLAG-CDC31* fragment and linearized pRS426-GPD vector were then assembled and the proper construct confirmed by DNA sequencing of the plasmid. pRS425-GFP-Spo20^51-91^ was constructed by cloning an EcoRI-XhoI fragment carrying the GFP fusion from pRS424-GFP-Spo20^51-91^ [31] into pRS425-TEF [50]. The plasmid pRS-Y2-myc-Myo5 carrying the myc-tagged *MYO5* and pRS425-Cas9-SkHIS3-381 for targeting double strand breaks by CRISPR have been described elsewhere [45, 51].

**Table 3.** Plasmids used in this study.

| Plasmid | Relevant features | Source |
|---|---|---|
| p426-GPD-FLAG-CDC31 | 2μ, *URA3*, P<sub>GPD1</sub>-*FLAG-CDC31* | This Study |
| TEP36 | 2μ, *LEU2*, P<sub>CUP1</sub>-*FLAG-CDC31* | [49] |
| pRS426-GPD | 2μ, *URA3*, P<sub>GPD1</sub> | [50] |
| pRS425-GFP-Spo20<sup>51-91</sup> | 2μ, *LEU2*, P<sub>TEF2</sub>- *GFP-SPO20*<sup>51-91</sup> | This Study |
| pRS424-GFP-Spo20<sup>51-91</sup> | 2μ, *TRP1*, P<sub>TEF2</sub>- *GFP-SPO20*<sup>51-91</sup> | [31] |
| pRS425-TEF | 2μ, *LEU2*, P<sub>TEF2</sub> | [50] |
| pRS-Y2-myc-Myo5 | 2μ, *URA3, myc-MYO5* | [51] |
| pRS316-MMM1 | CEN, *URA3*, *MMM1* | [15] |
| pRS425-Cas9-SkHIS3-381 | 2μ, *LEU2*, *CAS9*, SkHIS3-targeted guide RNA | [41] |

### AlphaFold structural predictions

Vps13^1-1800^ and full length Cdc31 and Cmd1 were cofolded using AlphaFold2 multimer at the colab website (https://colab.research.google.com/github/sokrypton/ColabFold/blob/main/AlphaFold2.ipynb) or Alphafold3 at the alphafold website (https://alphafoldserver.com).

### Protein extractions and immunoprecipitation

Trichloroacetic acid (TCA) precipitation for total protein extraction was performed as previously described [41]. 5 mL of exponentially growing cells were pelleted and resuspended in 5 mL of 5% TCA and incubated at 4° C for 10 min. After centrifugation at 4000 RPM (2000 x g) for 5 min in a tabletop centrifuge, the pellets were resuspended in 1 mL of room temperature acetone, transferred to 2 mL screw cap tubes and centrifuged at 11,000 x g for 5 min.

Supernatant was discarded and the pellets left to air dry overnight. Pellets were resuspended in 200 µL lysis buffer (TUNES Buffer: 100 mM Tris at pH 7.2, 6 M urea, 10 mM EDTA, 1% SDS, 0.4 M NaCl, 10% glycerol and 50 mM N-ethylmaleimide) and then broken up thoroughly using a pipet tip. 200 µL of 0.5 mm diameter Zirconia beads (BioSpec cat. 11079105z) were added and the cells lysed using FastPrep-24 cell lysis machine (MP Biomedical) at speed 6.5 for three pulses of 45 s each. After bead beating, lysates were incubated in 150 µL of 2 X sodium dodecyl sulfate (SDS) sample buffer (100 mM Tris-HCl pH 6.8, 4% SDS, 20% Glycerol, 0.2 mg/mL Bromophenol blue, 0.72 M 2-Mercaptoethanol) at 50°C for 10 min and then spun at 11,000 x g for 5 min before loading onto an SDS-polyacrylamide gel.

For IPs using FLAG antibodies, cells were grown in synthetic complete medium lacking uracil or leucine, as appropriate, to an optical density of approximately 1 and 6.0 × 10⁸ cells were harvested. Cells were washed once in ice-cold water and resuspended in 400 µL of lysis buffer (50 mM HEPES, pH 7.5, 150 mM NaCl, 5 mM EDTA, 1% Triton, protease inhibitors (1 tablet per 5ml; Roche cat. 04693159001), 1 mM PMSF, and 40 µg/mL zymolyase 100T (Fisher cat. Z1004) and divided between two 2 ml screw-cap tubes per sample. 200 µL of zirconia beads were added and cells lysed using the FastPrep machine as described above. Lysates were clarified by centrifugation at 11,000 × g for 5 min at 4°C. 500 µL of dilution buffer (10 mM Tris/Cl, pH 7.5, 150 mM NaCl, 5 mM EDTA, protease inhibitors, 1 mM PMSF, and 40 µg/mL zymolyase 100T) was added to the lysates and clarified lysates were incubated with 50 µL of anti-FLAG M2 affinity gel (Millipore Sigma cat. A2220) at 4°C for 4 h on a rocking platform. After incubation, samples were centrifuged at 11,000 × g for 2 min at 4°C, beads were washed twice with 500 µL and once with 200 µL of ice-cold wash buffer (50 mM HEPES, pH 7.5, 150 mM NaCl, and 5 mM EDTA), with centrifugation at 11,000 × *g* for 2 min at 4°C after each wash. The immunoprecipitated samples were incubated for 10 min in 50 µL of 2× SDS sample buffer.

GFP IPs were performed using a different lysis buffer than the FLAG IPs (10 mM Tris/Cl, pH 7.5, 150 mM NaCl, 5 mM EDTA, 0.5% Nonidet™ P40 Substitute, protease inhibitors, 1 mM PMSF, and 40 µg/mL zymolyase 100T). Proteins were precipitated by incubation with 20 µL of GFP-Trap agarose beads (ChromoTek cat. gta-100) for 1 h at 4°C on a rocking platform. All spins involving the GFP beads were done at 2,500 × g for 2 min at 4°C.

For the serial IPs, the lysate supernatants were retained after incubation with the anti-FLAG M2 affinity gel and subsequent centrifugation were then incubated with anti-GFP affinity beads for 1 hr at 4°C.

### Western Blot Analysis

Proteins were fractionated by SDS-PAGE and transferred to polyvinylidene difluoride membranes (PVDF; Merck Millipore cat. IPVH85R), as described [41]. To detect Vps13 or Myo5, 7.5% gels were run at 100 volts for 2.5 h and transferred to PVDF at 50 mA overnight. To visualize Cmd1 or Cdc31, 12.5% gels were run at 200 volts for 30 mins and then transferred to PVDF at 300 mA for 1 hr. α-Vps13 antibodies [41] and α-myc antibodies (BioRad cat. MCA2200GA) were used at 1:500 dilution; α-GFP (Clontech Laboratories cat. 632381) and α-FLAG antibodies (Millipore Sigma cat. F3165) were used at 1:1000 dilution; α-Pgk1 anitbodies (Invitrogen cat. 495250) were used at 1:5000 dilution. Horseradish peroxidase (HRP)-conjugated goat anti-rabbit IgG and goat anti-mouse IgG (Invitrogen) secondary antibodies were used at 1:5,000 dilution. HRP activity was detected using a Clarity Western ECL Substrate kit (BioRad cat. 170-5060) and a BioRad Chemidoc machine.

### Mass spectrometry

To obtain cell pellets of cells in meiosis II, strains ADY200 (containing pRS425-GFP-Spo20^51-91^) and JSP445 (*vps13-GFP*) were sporulated synchronously using an estradiol-inducible Ndt80 [32]. First, cells were grown overnight in 10ml of SD-Leu or YPD medium, respectively. The following day, the cultures were diluted into 150 mL of YPA medium and grown overnight. On the third day, cells were washed with sterile distilled water and resuspended in 150 mL of 2% potassium acetate sporulation medium. After 6 h of incubation, β-estradiol was added to a final concentration of 1 μM from a 10 mM stock solution. Cells were incubated for an additional 3 hr, progression into meiosis II was confirmed by DAPI staining, and then cells were harvested by centrifugation. Cell pellets were stored at −80 °C until the GFP pull-down procedure.

GFP pull-downs were performed as previously described by [52]. Cell pellets were divided equally among four screw-cap microcentrifuge tubes, and each pellet was resuspended in 200 μL of lysis buffer containing 10 mM Tris/HCl (pH 7.5), 150 mM NaCl, 0.5 mM EDTA, and 0.5% Nonidet P-40, supplemented with protease inhibitors, 1 mM PMSF, and 10μg/mL zymolyase 100T. Zirconia beads (200 mg) were then added to each tube. Cells were lysed using a FastPrep-24 cell lysis machine at a speed setting of 6.5 for two 45-s pulses.

The lysates were centrifuged at 11,000 × g for 5 min at 4 °C, and the supernatants were collected into a new microcentrifuge tube. GFP-Trap agarose beads (30 μL) pre-washed with 500 μL of washing buffer containing 10 mM Tris/HCl (pH 7.5), 150 mM NaCl, and 0.5 mM EDTA, were added to the pooled supernatants. The mixture was incubated with rotation for 2 h at 4 °C. The agarose beads were then collected by centrifugation at 2,500 × g for 5 min at 4 °C and washed twice with 500 μL of washing buffer. The washed beads were collected and used for mass spectrometry analysis. Peptides were released from the beads by trypsin digestion overnight at 37° C. After digestion the beads were collected by centrifugation and the supernatants containing the peptides were analysed by mass spectrometry.

Liquid chromatography tandem mass spectrometry (LC-MS/MS) analysis was performed on a Thermo Scientific LTQ ion-trap mass spectrometer equipped with an electrospray ionization (ESI) source. Peptides were chromatographically separated using a multistep, three-solvent liquid chromatography gradient as previously described [53]. Samples were maintained at 4 °C in the autosampler, and 10 µL was injected for analysis. The total chromatographic run time was 185 min.

The mass spectrometer was operated in positive-ion mode with the following source parameters: Spray Voltage, 2.0 kV; Capillary Temperature, 200 °C; Tube lens voltage, 100 V. Full-MS scans were acquired from 400–2000 m/z with an automatic gain control (AGC) target of 30,000 ions, a maximum ion injection time of 50 ms, and one microscan.

Data dependent acquisition was performed by selecting the five most intense precursor ions from each full-MS scan for collision-induced dissociation (CID) with a normalized collision energy of 35%, activation q of 0.25, and activation time of 30 ms. A 3.0 m/z precursor isolation width was applied along with a minimum signal intensity of 500, AGC target of 10,000 ions, a maximum ion injection time of 100 ms, and one microscan.

Dynamic exclusion was enabled with a repeat count of 3 within 30 s, an exclusion duration of 180 s, an exclusion list size of 500, and an exclusion mass window of ±1.5 m/z. Charge-state screening and charge-state rejection were disabled. Samples were maintained at 4 °C in the autosampler, and 10 µL was injected for analysis. The total MS acquisition time was 185 minutes.

To identify candidate Vps13 interactors, proteins identified in the precipitates from ADY200 were removed from the list of proteins co-precipitating with VPS13-GFP, as were ribosomal proteins and proteins known to localize to the mitochondrial matrix or extracellularly. The remaining candidates were examined for the number of unique peptides assigned to the protein as well as the percentage of the protein sequence covered by the identified peptides. The list in Table 1 is based on cutoffs of 2 unique peptides and greater than 10% coverage.

### Phenotypic assays

Assays for CPY secretion, ER-Phagy, and sporulation were performed as described previously [41]. For the CPY assay, 4-fold serial dilutions of exponentially growing cultures (starting at 1 × 10^7^ cells/mL) were spotted onto SC plates. Nitrocellulose membranes were placed on top of the cells, and the plates were incubated overnight. Membranes were then washed with Western blot washing buffer (10 mM Tris-HCl pH 7.5, 2.5 mM EDTA pH 8.0, 50 mM NaCl, 0.05% Tween-20) 5 times for five minutes each to remove cells and probed with monoclonal anti-CPY antibodies (Life Technologies cat. A6428) at 1:2000 dilution, followed by the HRP conjugated goat anti-mouse IgG secondary antibody (Invitrogen, cat. 31430) at 1:5000 dilution.

To examine ER-phagy, rapamycin was added (final concentration of 200 ng/mL) to exponentially growing cultures of cells expressing the peripheral ER marker Rtn1-GFP and the cultures were incubated overnight. Cells were then examined for the presence GFP fluorescence in the vacuole by epifluorescence microscopy.

To assay sporulation, patches of diploid strains were incubated on SPO medium at 30° for 24 hours before being transferred to slides and examined for the presence of spores by light microscopy. Six independent diploids of each genotype were sporulated and at least 400 cells were counted in each culture for the sporulation efficiency measurements.

To assay synthetic interactions with *mmm1*Δ, exponentially growing cultures of BY4741, KO1, KO4, NDY12, and NDY24 were adjusted to 1 × 10⁷ cells/mL based on their optical density at 660 nm, 10-fold serial dilutions were spotted onto YPD medium, and plates incubated at 30°C and 37°C for three days.

### Microscopy

All imaging for the ER-phagy and sporulation assays as well as Vps13 localization was performed on a Zeiss Imager.Z2 microscope (Carl Zeiss) with a Zeiss Axiocam 702 mono digital camera. ZEN 3.0 (Blue edition) software was used to acquire images.

## Acknowledgements

The authors thank Sai Zhou and Aviva Diamond for strains; Hiten Madhani for the GFP fusion collection; Liza Pon, Hideki Nakanishi, and Kiran Madura for plasmids; Kai Zhang for help with data analysis; and members of the Neiman lab for advice over the course of these studies. We thank Nancy Hollingsworth for comments on the manuscript. This work was supported by NIH R01 grants GM072540 and GM145606 to A.M.N.

## Literature Cited

1. Braschi B, Bruford EA, Cavanagh AT, Neuman SD, Bashirullah A. The bridge-like lipid transfer protein (BLTP) gene group: introducing new nomenclature based on structural homology indicating shared function. Hum Genomics. 2022;16(1):66. Epub 2022/12/03. doi: 10.1186/s40246-022-00439-3. PubMed PMID: 36461115

2. Neuman SD, Levine TP, Bashirullah A. A novel superfamily of bridge-like lipid transfer proteins. Trends in Cell Biology. 2022;32(11):962–74. Epub 2022/05/02. doi: 10.1016/j.tcb.2022.03.011. PubMed PMID: 35491307.

3. Levine TP. Sequence Analysis and Structural Predictions of Lipid Transfer Bridges in the Repeating Beta Groove (RBG) Superfamily Reveal Past and Present Domain Variations Affecting Form, Function and Interactions of VPS13, ATG2, SHIP164, Hobbit and Tweek. Contact. 2022;5:251525642211343. Epub 2022/12/27. doi: 10.1177/25152564221134328. PubMed PMID: 36571082.

4. Hanna M, Guillen-Samander A, De Camilli P. RBG Motif Bridge-Like Lipid Transport Proteins: Structure, Functions, and Open Questions. Annual Review of Cell and Developmental Biology. 2023. Epub 2023/07/05. doi: 10.1146/annurev-cellbio-120420-014634. PubMed PMID: 37406299.

5. Kolehmainen J, Black GC, Saarinen A, Chandler K, Clayton-Smith J, Traskelin AL, et al. Cohen syndrome is caused by mutations in a novel gene, *COH1*, encoding a transmembrane protein with a presumed role in vesicle-mediated sorting and intracellular protein transport. American Journal of Human Genetics. 2003;72(6):1359–69. PubMed PMID: 12730828.

6. Lesage S, Drouet V, Majounie E, Deramecourt V, Jacoupy M, Nicolas A, et al. Loss of *VPS13C* Function in Autosomal-Recessive Parkinsonism Causes Mitochondrial Dysfunction and Increases PINK1/Parkin-Dependent Mitophagy. American Journal of Human Genetics. 2016;98(3):500–13. doi: 10.1016/j.ajhg.2016.01.014. PubMed PMID: 26942284.

7. Rampoldi L, Dobson-Stone C, Rubio JP, Danek A, Chalmers RM, Wood NW, et al. A conserved sorting-associated protein is mutant in chorea-acanthocytosis. Nature Genetics. 2001;28(2):119–20. PubMed PMID: 11381253.

8. Seong E, Insolera R, Dulovic M, Kamsteeg EJ, Trinh J, Bruggemann N, et al. Mutations in *VPS13D* lead to a new recessive ataxia with spasticity and mitochondrial defects. Annals of Neurology. 2018;83(6):1075–88. doi: 10.1002/ana.25220. PubMed PMID: 29604224

9. Swan LE. VPS13 and bridge-like lipid transporters, mechanisms, and mysteries. Front Neurosci. 2025;19:1534061. Epub 2025/05/13. doi: 10.3389/fnins.2025.1534061. PubMed PMID: 40356703.

10. Dziurdzik SK, Conibear E. The Vps13 Family of Lipid Transporters and Its Role at Membrane Contact Sites. Int J Mol Sci. 2021;22(6). Epub 2021/04/04. doi: 10.3390/ijms22062905. PubMed PMID: 33809364.

11. Velayos-Baeza A, Vettori A, Copley RR, Dobson-Stone C, Monaco AP. Analysis of the human *VPS13* gene family. Genomics. 2004;84(3):536–49. PubMed PMID: 15498460.

12. Bankaitis VA, Johnson LM, Emr SD. Isolation of yeast mutants defective in protein targeting to the vacuole. Proceedings of the National Academy of Sciences of the United States of America. 1986;83(23):9075–9. PubMed PMID: 3538017.

13. Chen S, Mari M, Parashar S, Liu D, Cui Y, Reggiori F, et al. Vps13 is required for the packaging of the ER into autophagosomes during ER-phagy. Proceedings of the National Academy of Sciences of the United States of America. 2020;117(31):18530–9. Epub 2020/07/22. doi: 10.1073/pnas.2008923117. PubMed PMID: 32690699.

14. Park JS, Neiman AM. *VPS13* regulates membrane morphogenesis during sporulation in *Saccharomyces cerevisiae*. Journal of Cell Science. 2012;125:3004–11. PubMed PMID: 22442115.

15. Park JS, Thorsness MK, Policastro R, McGoldrick L, Hollingsworth NM, Thorsness PE, et al. Yeast Vps13 promotes mitochondrial function and is localized at membrane contact sites. Molecular Biology of the Cell. 2016;27:2435–49.

16. Lang AB, John Peter AT, Walter P, Kornmann B. ER-mitochondrial junctions can be bypassed by dominant mutations in the endosomal protein Vps13. The Journal of Cell Biology. 2015;210(6):883–90. doi: 10.1083/jcb.201502105. PubMed PMID: 26370498.

17. Bean BDM, Dziurdzik SK, Kolehmainen KL, Fowler CMS, Kwong WK, Grad LI, et al. Competitive organelle-specific adaptors recruit Vps13 to membrane contact sites. The Journal of Cell Biology. 2018;217(10):3593–607. doi: 10.1083/jcb.201804111. PubMed PMID: 30018089.

18. John Peter AT, Herrmann B, Antunes D, Rapaport D, Dimmer KS, Kornmann B. Vps13-Mcp1 interact at vacuole-mitochondria interfaces and bypass ER-mitochondria contact sites. The Journal of Cell Biology. 2017;216(10):3219–29. doi: 10.1083/jcb.201610055. PubMed PMID: 28864540.

19. Park JS, Okumura Y, Tachikawa H, Neiman AM. *SPO71* encodes a developmental stage-specific partner for *VPS13* in *Saccharomcyes cerevisiae*. Eukaryotic Cell. 2013;12:1530– 7. PubMed PMID: 24036347.

20. Kumar N, Leonzino M, Hancock-Cerutti W, Horenkamp FA, Li P, Lees JA, et al. VPS13A and VPS13C are lipid transport proteins differentially localized at ER contact sites. The Journal of Cell Biology. 2018;217(10):3625–39. doi: 10.1083/jcb.201807019. PubMed PMID: 30093493.

21. Yeshaw WM, van der Zwaag M, Pinto F, Lahaye LL, Faber AIE, Gomez-Sanchez R, et al. Human VPS13A is associated with multiple organelles and influences mitochondrial morphology and lipid droplet motility. Elife. 2019;8. doi: ARTN e43561 10.7554/eLife.43561. PubMed PMID: WOS:000459683300001.

22. De M, Oleskie AN, Ayyash M, Dutta S, Mancour L, Abazeed ME, et al. The Vps13p-Cdc31p complex is directly required for TGN late endosome transport and TGN homotypic fusion. The Journal of Cell Biology. 2017;216(2):425–39. doi: 10.1083/jcb.201606078. PubMed PMID: 28122955.

23. Kilmartin JV. Sfi1p has conserved centrin-binding sites and an essential function in budding yeast spindle pole body duplication. The Journal of Cell Biology. 2003;162(7):1211–21. Epub 20030922. doi: 10.1083/jcb.200307064. PubMed PMID: 14504268.

24. Hartwell LH, Mortimer RK, Culotti J, Culotti M. Genetic Control of the Cell Division Cycle in Yeast: V. Genetic Analysis of cdc Mutants. Genetics. 1973;74(2):267–86. doi: 10.1093/genetics/74.2.267. PubMed PMID: 17248617.

25. Baum P, Furlong C, Byers B. Yeast gene required for spindle pole body duplication: homology of its product with Ca2+-binding proteins. Proceedings of the National Academy of Sciences of the United States of America. 1986;83(15):5512–6. PubMed PMID: 3526331.

26. Fischer T, Rodriguez-Navarro S, Pereira G, Racz A, Schiebel E, Hurt E. Yeast centrin Cdc31 is linked to the nuclear mRNA export machinery. Nature Cell Biology. 2004;6(9):840–8. Epub 20040815. doi: 10.1038/ncb1163. PubMed PMID: 15311284.

27. Spang A, Courtney I, Fackler U, Matzner M, Schiebel E. The calcium-binding protein cell division cycle 31 of *Saccharomyces cerevisiae* is a component of the half bridge of the spindle pole body. The Journal of Cell Biology. 1993;123(2):405–16. doi: 10.1083/jcb.123.2.405. PubMed PMID: 8408222.

28. Michaelis AC, Brunner AD, Zwiebel M, Meier F, Strauss MT, Bludau I, et al. The social and structural architecture of the yeast protein interactome. Nature. 2023;624(7990):192–200. Epub 20231115. doi: 10.1038/s41586-023-06739-5. PubMed PMID: 37968396.

29. Li D, Wang X, Hao H, Eden J, Hu B, Walsh EE, et al. Cryo-EM structure of soluble VPS13C suggests its regulation by a conformational switch and by calmodulin. Molecular Cell. 2026;86(14):2843–57 e10. Epub 20260707. doi: 10.1016/j.molcel.2026.06.028. PubMed PMID: 42413490.

30. Evans R, O’Neill M, Pritzel A, Antropova N, Senior AW, Green T, et al. Protein complex prediction with AlphaFold-Multimer. bioRxiv. 2022. doi: 10.1101/2021.10.04.463034.

31. Nakanishi H, de los Santos P, Neiman AM. Positive and negative regulation of a SNARE protein by control of intracellular localization. Molecular Biology of the Cell. 2004;15(4):1802–15. PubMed PMID: 14742704.

32. Benjamin KR, Zhang C, Shokat KM, Herskowitz I. Control of landmark events in meiosis by the CDK Cdc28 and the meiosis-specific kinase Ime2. Genes & Development. 2003;17(12):1524–39. PubMed PMID: 12783856.

33. Carlile TM, Amon A. Meiosis I is established through division-specific translational control of a cyclin. Cell. 2008;133(2):280–91. PubMed PMID: 18423199.

34. Neiman AM. Prospore membrane formation defines a developmentally regulated branch of the secretory pathway in yeast. The Journal of Cell Biology. 1998;140(1):29–37. PubMed PMID: 9425151.

35. Davis TN, Urdea MS, Masiarz FR, Thorner J. Isolation of the yeast calmodulin gene: calmodulin is an essential protein. Cell. 1986;47(3):423–31. doi: 10.1016/0092-8674(86)90599-4. PubMed PMID: 3533275.

36. van Heusden GP, Griffiths DJ, Ford JC, Chin AWTF, Schrader PA, Carr AM, et al. The 14-3-3 proteins encoded by the *BMH1* and *BMH2* genes are essential in the yeast *Saccharomyces cerevisiae* and can be replaced by a plant homologue. European Journal of Biochemistry / FEBS. 1995;229(1):45–53. PubMed PMID: 7744048.

37. Segal D, Maier S, Mastromarco GJ, Qian WW, Nabeel-Shah S, Lee H, et al. A central chaperone-like role for 14-3-3 proteins in human cells. Molecular Cell. 2023;83(6):974–93 e15. doi: 10.1016/j.molcel.2023.02.018. PubMed PMID: 36931259.

38. Bers DM, Patton CW, Nuccitelli R. A practical guide to the preparation of Ca(2+) buffers. Methods Cell Biol. 2010;99:1–26. doi: 10.1016/B978-0-12-374841-6.00001-3. PubMed PMID: 21035681.

39. Geli MI, Wesp A, Riezman H. Distinct functions of calmodulin are required for the uptake step of receptor-mediated endocytosis in yeast: the type I myosin Myo5p is one of the calmodulin targets. EMBO J. 1998;17(3):635–47. doi: 10.1093/emboj/17.3.635. PubMed PMID: 9450989.

40. Roberts CJ, Raymond CK, Yamashiro CT, Stevens TH. Methods for studying the yeast vacuole. Methods Enzymol. 1991;194:644–61. Epub 1991/01/01. doi: 10.1016/0076-6879(91)94047-g. PubMed PMID: 1706462.

41. Park JS, Hollingsworth NM, Neiman AM. Genetic dissection of Vps13 regulation in yeast using human disease mutations. Int J Mol Sci. 2021;22:6200.

42. Smolders S, Philtjens S, Crosiers D, Sieben A, Hens E, Heeman B, et al. Contribution of rare homozygous and compound heterozygous VPS13C missense mutations to dementia with Lewy bodies and Parkinson’s disease. Acta Neuropathol Commun. 2021;9(1):25. Epub 2021/02/14. doi: 10.1186/s40478-021-01121-w. PubMed PMID: 33579389.

43. Weiss B, Dunham M, Gartenberg MR. Methods in Yeast Genetics and Genomics. Cold Spring Harbor, NY: Cold Spring Harbor Laboratory Press; 2015.

44. Longtine MS, McKenzie A, 3rd, Demarini DJ, Shah NG, Wach A, Brachat A, et al. Additional modules for versatile and economical PCR-based gene deletion and modification in Saccharomyces cerevisiae. Yeast. 1998;14(10):953–61. PubMed PMID: 9717241.

45. Jin L, Zhang K, Xu Y, Sternglanz R, Neiman AM. Sequestration of mRNAs modulates the timing of translation during meiosis in budding yeast. Molecular and Cellular Biology. 2015;35(20):3448–58. doi: 10.1128/MCB.00189-15. PubMed PMID: 26217015.

46. Huh WK, Falvo JV, Gerke LC, Carroll AS, Howson RW, Weissman JS, et al. Global analysis of protein localization in budding yeast. Nature. 2003;425(6959):686–91. PubMed PMID: 14562095.

47. Berchowitz LE, Gajadhar AS, van Werven FJ, De Rosa AA, Samoylova ML, Brar GA, et al. A developmentally regulated translational control pathway establishes the meiotic chromosome segregation pattern. Genes & Development. 2013;27(19):2147–63. Epub 2013/10/12. doi: 10.1101/gad.224253.113. PubMed PMID: 24115771.

48. Diamond A, Park JS, Inoue I, Tachikawa H, Neiman AM. The APC targeting subunit Ama1 links meiotic exit to cytokinesis during sporulation in *Saccharomyces cerevisiae*. Molecular Biology of the Cell. 2008;20:134–45. PubMed PMID: 18946082.

49. Chen L, Madura K. Centrin/Cdc31 is a novel regulator of protein degradation. Molecular and cellular biology. 2008;28(5):1829–40. Epub 20071226. doi: 10.1128/MCB.01256-07. PubMed PMID: 18160718.

50. Mumberg D, Muller R, Funk M. Yeast vectors for the controlled expression of heterologous proteins in different genetic backgrounds. Gene. 1995;156(1):119–22. PubMed PMID: 7737504.

51. Anderson BL, Boldogh I, Evangelista M, Boone C, Greene LA, Pon LA. The Src homology domain 3 (SH3) of a yeast type I myosin, Myo5p, binds to verprolin and is required for targeting to sites of actin polarization. The Journal of Cell Biology. 1998;141(6):1357–70. doi: 10.1083/jcb.141.6.1357. PubMed PMID: 9628892.

52. Park JS, Hu Y, Hollingsworth NM, Miltenberger-Miltenyi G, Neiman AM. Interaction between VPS13A and the XK scramblase is important for VPS13A function in humans. Journal of Cell Science. 2022;135(17). Epub 2022/08/12. doi: 10.1242/jcs.260227. PubMed PMID: 35950506.

53. Abraham AB, Bronstein R, Chen EI, Koller A, Ronfani L, Maletic-Savatic M, et al. Members of the high mobility group B protein family are dynamically expressed in embryonic neural stem cells. Proteome Sci. 2013;11(1):18. Epub 20130427. doi: 10.1186/1477-5956-11-18. PubMed PMID: 23621913.

54. Winzeler EA, Shoemaker DD, Astromoff A, Liang H, Anderson K, Andre B, et al. Functional characterization of the *S. cerevisiae* genome by gene deletion and parallel analysis. Science. 1999;285(5429):901–6. PubMed PMID: 10436161.

55. Neiman AM, Katz L, Brennwald PJ. Identification of domains required for developmentally regulated SNARE function in *Saccharomyces cerevisiae*. Genetics. 2000;155(4):1643–55. PubMed PMID: 10924463.

56. Nakanishi H, Suda Y, Neiman AM. Erv14 family cargo receptors are necessary for ER exit during sporulation in *Saccharomyces cerevisiae*. Journal of Cell Science. 2007;120(Pt 5):908–16. PubMed PMID: 17298976.

